# Transgenerational inheritance of shuffled symbiont communities in the coral *Montipora digitata*

**DOI:** 10.1101/264796

**Authors:** Kate M. Quigley, Bette L. Willis, Carly D. Kenkel

## Abstract

Adult organisms may “prime” their offspring for environmental change through a number of genetic and non-genetic mechanisms, termed parental effects. Some coral species can alter their thermal tolerance by shuffling the proportions of *Symbiodinium* types within their endosymbiotic communities, but it is unclear if this plasticity can be transferred to offspring in corals that have maternal symbiont transmission. We evaluated symbiont community composition in tagged colonies of *Montipora digitata* from Orpheus Island, Australia, over two successive annual spawning seasons, the second of which overlapped with the 2016 mass coral bleaching event on the Great Barrier Reef. We applied amplicon sequencing of the ITS2 locus to samples of four families (parent colonies and 10-12 eggs per family) to characterize their potential for symbiont shuffling and to determine if shuffled abundances were preserved in gametes. Symbiont cell densities and photochemical efficiencies of the symbionts’ photosystem II differed significantly among adults in 2016, suggesting differential responses to increased temperatures. Although abundances of the dominant symbiont haplotype, a representative of clade C15, did not differ among families or over time, low-abundance (“background”) ITS2 types differed more among years (2015 vs. 2016) than between life stages (parent vs. offspring). Results indicate that background symbiont shuffling can occur in a canonically ‘stable’ symbiosis, and that such plastic changes to the symbiont community are heritable. To our knowledge, this is the first evidence that shuffled *Symbiodinium* communities can be inherited by early life-history stages and supports the hypothesis that plastic changes in microbial communities may serve as a mechanism of rapid coral acclimation to changing environmental conditions.

## Introduction

Acclimatization to fluctuating environmental conditions through phenotypic plasticity can promote the persistence of populations and may facilitate subsequent genetic rescue in an era of global climate change [1,2]. In some cases, plasticity may also be associated with transgenerational effects, whereby the performance of offspring is influenced by environmental conditions experienced by the parents [3]. Inheritance of altered parental phenotypes via transgenerational plasticity may further buffer populations by facilitating acclimatization over generations [4,5].

In corals, the Adaptive Bleaching Hypothesis (ABH) [6] has been proposed as a potential acclimatization mechanism that could protect adult corals from changing environmental conditions through alteration of their endosymbiotic *Symbiodinium* communities [7–10]. The proportional abundances of *Symbiodinium* types within adult coral tissues can vary (“shuffling”) to favour types that appear to be more tolerant of prevailing environmental conditions [10–14]. Such plastic changes to the *Symbiodinium* community have the potential to decrease the predicted frequency of bleaching, the breakdown of the host-symbiont relationship as a result of stress, by 14% [15]. However, it is unclear whether acclimatory changes to the *Symbiodinium* community are limited to the lifetime of an adult coral or if shuffled communities are inherited across generations in coral species capable of vertical symbiont transmission.

Vertically-transmitting coral species provision eggs or planula larvae with *Symbiodinium* types predominantly from the maternal parent [16]. Genetic factors influencing the overall composition of the *Symbiodinium* community in these early life stages are inherited in at least three species [17,18]. As a heritable trait, genetic constraints (e.g. host controlled immunity and recognition) may limit the capacity of vertical transmitters to modify the diversity of their symbiont communities that are not found in species with environmental acquisition [19]. Padilla-Gamiño *et al.* [20] examined *Symbiodinium* communities in adults and eggs of the vertically-transmitting coral *Montipora capitata* across years, and found that while community composition was predominantly faithfully transmitted, the community did not change over time. In contrast, *Porites astreoides* planulae differed in their symbiont communities compared to adults and juveniles [21]. However, transgenerational inheritance of other traits, including symbiont switching, has been suggested [22]. For example, larvae of the vertically-transmitting coral *Pocillopora damicornis* exhibited altered respiration rates when parents were exposed to elevated pCO_2_ during the development of brooded larvae [23]. Given the major role symbionts play in coral thermal tolerance [12], evaluations of the potential for increased ocean temperatures to alter maternal transmission of *Symbiodinium* and of the capacity for rapid acclimation and adaptation of offspring through transgenerational mechanisms are needed.

The importance of background (low-abundance) *Symbiodinium* types for the capacity of corals to alter their thermal tolerance is becoming more widely recognised, in light of growing evidence that the capacity to shuffle *Symbiodinium* usually involves a change in the abundance of background species [14,24–26]. *Symbiodinium* from clade D, the taxa most often reported as instrumental in post-bleaching recovery in corals, normally exist at low levels in coral hosts, even as low as 100 D cells to 10,000 host cells per cm^2^ [27,28].

Recently, Bay et al. (2016) found that some colonies of *Acropora millepora* from a naturally cooler population with a ratio of at least 3:1000 D:C type symbionts were able to shuffle to a D-dominated community during stress, and subsequently exhibited increased survival and recovery. Background symbiont types are also known to be functionally important in other host-microbe associations. For example, rare members of the bacterial community play disproportionally important roles in nitrogen cycling in their hosts (e.g. *Desulfosporosinus* in peat soil [29]). Therefore, changes in the relative abundances of background *Symbiodinium* types have the potential to change the host phenotype, thereby contributing to its stress tolerance and recovery [24,25].

We took advantage of a natural thermal stress event, which caused severe bleaching in corals on the Great Barrier Reef (GBR) in 2016 [30], to evaluate the potential for transgenerational inheritance of shuffled symbiont communities in a vertically-transmitting coral. Using amplicon sequencing of the ITS2 locus, we compared *Symbiodinium* communities in a sample set comprised of four colonies of *Montipora digitata* and 10-12 eggs from each colony collected during the 2016 mass bleaching event to a replicate sample set collected during the previous summer (2015; no thermal anomaly). Although this species is typically dominated by C15-type symbionts, it has been shown to host background Clade D and A types [17]. We evaluated the null hypothesis that symbiont communities in *M. digitata* are stable in response to thermal stress and highly heritable ([17], H_0_, figure 1A) and three possible alternative hypotheses: H_1_- symbiont community composition is stable, but not highly heritable (no shuffling, no parental effects); H_2_- symbiont community composition is plastic, and highly heritable (shuffling and parental effects); H3- symbiont community composition is plastic, but not heritable (shuffling, no parental effects).

**Figure 1.**
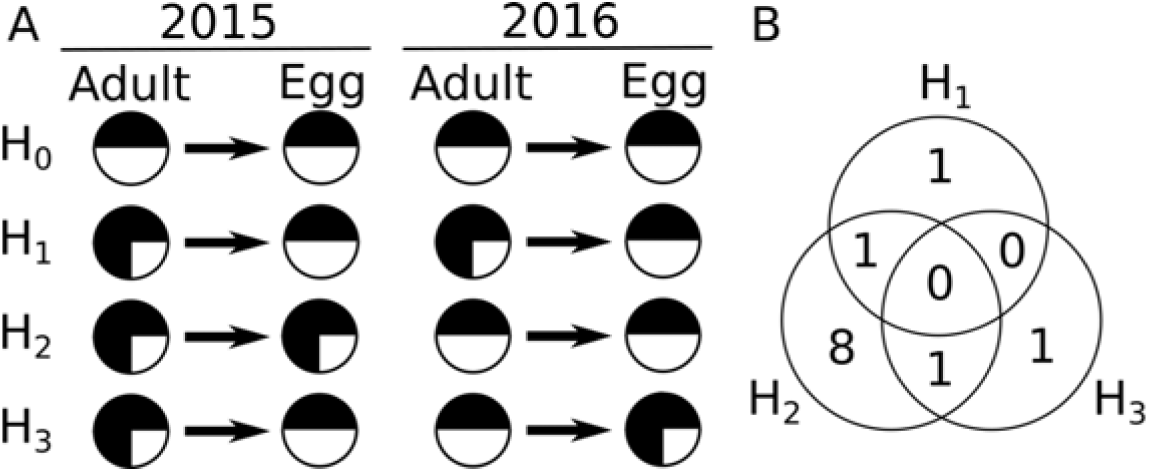
(A) Schematic of hypotheses for potential parental effects and/or shuffling across sampling years. (B) Venn diagram showing the number of significantly differentially abundant (P_adj_ < 0.05) sequence variants by hypothesis.

## Materials and methods

### Coral spawning and sample collection

Thirty-two colonies of *Montipora digitata* were collected from Hazard and Pioneer Bays in the Palm Island group on the Great Barrier Reef (GBR) on the 30^th^ of March and 1^st^ of April 2015. The same colonies were re-collected from Hazard Bay in April 2016 three days before the full moon, with the exception of colonies 4, 5, 8, 15 and 25, which were presumed dead (n_2016_ =27). In both years, colonies were placed in constant-flow 0.5 μM (2015) or 0.25 μM (2016) filtered seawater in outdoor raceways at Orpheus Island Research Station. Egg-sperm bundles were collected from nine colonies in 2015 (23^rd^ April) and four colonies in 2016 (24^th^ April). Eggs were filtered through a 100 μm mesh and rinsed three times to remove sperm and then individually preserved in 100% ethanol. Branches (n_2015_ = 1 branch/colony; n_2016_ = 3 branches/colony) were in 100% ethanol for genetic analysis of the *Symbiodinium* communities.

### Evaluating adult physiology during spawning in 2016 (El Niño year)

To determine the impacts of the El Niño-driven mass thermal stress event on *M. digitata* in 2016, three traits commonly used to evaluate holobiont health were measured: effective quantum yield of photosystem II (YII), colony coloration, and *Symbiodinium* cell density. YII was measured using Pulse Amplitude Modulated fluorometry (PAM) with a fibre optic cable (diving-PAM, Waltz). PAM measurements were taken at mid-day (~12:30) one day prior to spawning (22^nd^) and two days during the spawning period (23^rd^, 24^th^). Three measurements were taken per colony at each time point, with the instrument set at an Intensity of 12, Gain of 4 and F0 between 1.5 - 3. YII represents the percent quanta used by photosystem II of the *Symbiodinium* cells and is an indication of photosystem functioning and stress during light reactions. To test for significant differences in YII due to colony identity or day (as categorical factors), linear mixed models were fit using the package ‘nlme’ in R [31]. Replicate fluorescence measurements per colony were set as a random effect and uniform compound symmetry correlation structure (CorCompSymm) with Day as input values were used to account for autocorrelation issues. Analysis of Deviance Type II tests from the ‘car’ package were used to determine overall main and interactive effects [32]. The ‘glht’ package was used to extract Tukey p-adjusted values for multiple comparisons from linear models to determine if YII values differed among colonies [33]. Linear models with the same structure were used to test for significant differences in YII values between spawning and non-spawning colonies. Assumptions of homogeneity of variance, linearity, normality and autocorrelation were checked and tests modified where appropriate.

The presence of bleaching at the time of collection was assessed by photographing each colony alongside the CoralWatch color chart [34] to calibrate light conditions for image analysis. Three replicate RGB measurements were taken across each branch in ImageJ, resulting in continuous numerical values between 2.17 and 6, where values greater than 6 were designated as 6 [34]. The average of RGB color at each of the three randomly selected points per image were calibrated using photo-specific linear equations determined from analyses of the simultaneously photographed CoralWatch color chart. A negative binomial model (glmer function from ‘lme4’) was used to assess if mean bleaching status varied significantly with colony identity (colony = fixed effect, replicate color chart measures per colony = random effect). No deviations in homogeneity of variance, zero-inflation or over dispersion were detected. Likelihood ratio tests (LRT) showed that there was no significant effect of spawning status (spawning or not) or its interaction with dam identity on bleaching status (both Pr (>Chisq) = 1), thus these terms were dropped. LRT and effect sizes were used to assess the overall effect of colony identity on bleaching status (if the means differed significantly across colonies) by comparing the full model to the null model without colony identity using p-values calculated from a chi-squared distribution. *Symbiodinium* density was determined from spawning colonies using a Neubauer Hemocytometer (Optik Labor, UK). Triplicate cell counts were performed after nubbins were tissue blasted and standardized to the surface area of each nubbin. Surface areas were calculated using the wax dipping method, using 11 reference standard cylinders [35].

#### Symbiodinium community composition analysis

*Symbiodinium* communities in adults and eggs in both 2015 and 2016 were genotyped using paired-end Illumina Miseq amplicon sequencing (2 × 250bp) of the ITS2 locus [36] at the Genome Sequencing and Analysis Facility at the University of Texas at Austin (USA). Samples from all spawning adults (n_2015_ = 9 branches, 1 branch per coral; n_2016_ = 12 branches, 3 branches from each of 4 corals) and 10-12 eggs from each colony were sequenced in two independent runs (corresponding to 2015 versus 2016 samples).

Raw fastq reads were pre-filtered by removing any pairs that contained Illumina sequencing adapters (reads with an exact match of at least 12bp) or did not begin with the ITS2 amplicon primer sequence (reads lacking an exact match of at least 10bp), using BBDuk from the BBMap package version 37.75 (http://sourceforge.net/projects/bbmap/). ITS2 sequence variants were then inferred using the DADA2 pipeline [37] in R (v 3.4.1)[38]. The following analysis was completed twice, once for the full dataset, and once for a dataset in which the 2016 reads were randomly subsampled using seqtk (https://github.com/lh3/seqtk) to the mean abundance for the 2015 dataset post-quality filtering (7,000 PE reads per sample). Briefly, filtered fastq files were imported into R and read quality profiles were visually inspected. Reads were then filtered further, removing reads exhibiting matches to the phiX genome, reads with uncharacterized bases or reads with more than one expected error [37]. For reads passing these filters, ITS2 amplicon primers were trimmed prior to variant analysis [37].

Error rate models for forward and reverse reads were run until convergence, and estimated rates were visually inspected to assess fit with expected error rates. Sequence variants were inferred from the entire sequencing dataset using the default options for the “dada” command, which accounts for substitution and indel errors based on learned models, but the “BAND_SIZE” flag was set to 32 as is recommended for ITS data [37]. Inferred variants were further culled for length (294-304bp accepted) to remove potential products of non-specific priming, and chimeric sequences were removed. These high confidence sequence variants were taxonomically classified through a blast search against the GeoSymbio ITS2 database [39], and the best match was recorded. In cases where variants matched equally well to multiple references, all top hits were reported.

The MCMC.OTU package [40] was used to remove sample outliers with low counts overall (z-score <-2.5) and sequence variants that appeared in less than three unique samples prior to statistical analysis. Only the subset of families for which spawning was captured in the two consecutive years was retained in order to evaluate the impact of life stage and year on symbiont community composition.

A principal coordinates analysis (PCoA) was performed by computing Manhattan distances of log-transformed normalized variant counts. The resulting plots were visualized using the Phyloseq package [41] to examine the effects of the factors family, year and life stage on *Symbiodinium* community composition. The DESeq package [42] was used to construct a series of generalized linear models to evaluate differences in the abundance of sequence variants with respect to life stage (adult/egg) and year (2015/2016), including family (7, 9, 11, 24) as a blocking factor. Models were run for 30 iterations and, for models that did not converge, p-values were converted to NAs prior to applying a multiple test correction [43]. The Phyloseq package was used to plot results.

To assess the relationship between shuffling and bleaching, the composition of the *Symbiodinium* community was converted to a single quantitative metric (full description and methods in [17]) and correlated to the bleaching score. To facilitate comparisons between the two years, only samples collected from the centre of each colony in 2016 were used, in order to correspond to samples collected similarly in 2015.

Sequence variants of interest were aligned using Clustal Omega [44], as implemented on the EMBL-EBI web server (https://www.ebi.ac.uk/Tools/msa/clustalo/) and haplotype networks were visualized using the plot functions from the Pegas package [45]. Phylogenetic relationships among significant clade C-type OTUs were determined using the Phangorn package [46]. The Hasegawa-Kishino-Yano model of nucleotide substitution including invariant sites [47] was identified as the best fit model based on AIC, and was used to infer relationships among variants using 100 bootstrap replicates.

## Results

### *Montipora digitata* physiology during spawning in a severe bleaching year (2016)

Twelve of the experimental colonies showed variable signs of bleaching in 2016, as indicated by the bleaching status metric ranging from 3.08 ± 0.5 (colony 16) to 5.84 ± 0.2 (colony 10). Thirteen colonies did not exhibit any signs of bleaching or loss of pigmentation (bleaching status = 6) (figure 2A). Regardless of this variability in bleaching status across colonies, there was no significant relationship between bleaching status and colony identity, spawning status of the colony, or an interaction between the two (identity, spawning status, identity*spawning status: all NB LRT Pr(>Chisq) > 0.9996). *Symbiodinium* density varied significantly among the four colonies that spawned (LRT, df = 6, *p =* 0.0097) (figure 2 inset); with three spawning colonies exhibiting either no or light paling (5.8 ± 0.2) and colony 7 only moderate paling (4.8 ± 0.5).

**Figure 2.**
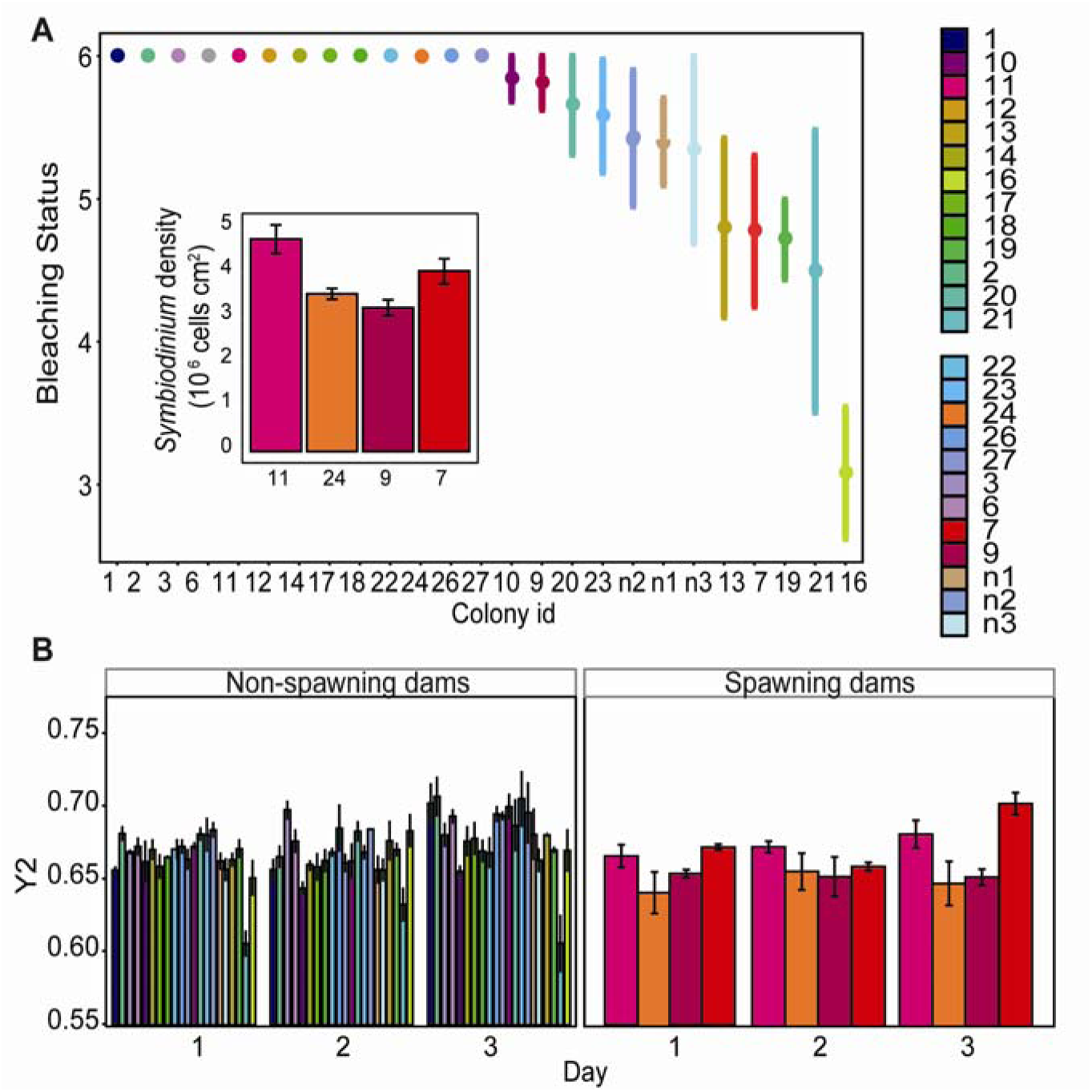
*Montipora digitata* physiological measures during bleaching. (A) Bleaching scores per colony during spawning 2016 (inset: average ± SE *Symbiodinium* density for spawning colonies). (B) Effective quantum yield of photosystem II (YII) during the spawning period for non-spawning and spawning colonies.

Overall, values for the effective quantum yield of photosystem II (YII) ranged from 0.699 ± 0.001 (colony n2) down to 0.618 ± 0.009 (colony 21). In general, YII values varied significantly by day, increasing from an average of 0.67 ± 0.002 to 0.69 ± 0.004 in the three days preceding and during spawning (n=25 colonies with three replicates measured per day); Likelihood Ratio Test-LRT, df = 2, *p =* 1.5e-07, figure 2B). Average YII measures also varied significantly among dams (LRT, df = 24, *p =* 2.2e-16), but in a consistent manner through time (no significant interaction between dam identity and day: LRT, df = 48, *p =* 0.127). Spawning colonies had significantly lower YII values compared to non-spawning colonies (0.66 ± 0.02 versus 0.68 ± 0.03; LME, df = 221, *p =* 1e-04).

### *Symbiodinium* community composition

On average, more reads per sample were obtained for the 2016 sample set than for the 2015 sample set (32,708 versus 14,020 2 × 250bp PE reads, respectively), and more reads were retained for the 2016 dataset following quality filtering (78% vs 33%). This resulted in an average of 5,138 reads per sample in 2015 and 26,566 reads per sample in 2016 being available for identification of ITS2 sequence variants (∼types).

A total of 34 high confidence types (those occurring in more than three unique samples) were identified from a subset of 107 high-read abundance samples (z-score > −2.5), representing the four families originating from colonies that spawned in both years (colonies/families 7, 9, 11 and 24). Of these 34 types, a single variant comprised 66% of the normalized read data across samples; a blast search against the GeoSymbio database (Franklin et al. 2012) identified it as a best match to *Symbiodinium* type C15. Additional background types, ranging in abundance from 17% to <1% of the normalized read data across counts also exhibited best matches to type C15 (26/34) or some derivative thereof (2/34 C15.h, 1/34 C15.6, 1/34 C15.8, 1/34 C15.9). The remaining three background types exhibited best matches to the C1 reference (comprising 0.05% of normalized reads) and two Clade D variants (best-matches to D1 and D1a, comprising 0.04% and 0.014% of normalized reads, respectively).

Principal coordinates analysis revealed that year was the main factor differentiating symbiont communities among samples (figure 3). This pattern could have been driven by the greater quantity of read data generated for the 2016 dataset. Therefore, the analysis was repeated on a dataset in which high coverage samples from 2016 were randomly subsampled to 7,000 PE reads to mimic a reduced sequencing effort. The same major sequence variants were recovered and year remained the main factor differentiating samples (figure S1,2).

**Figure 3.**
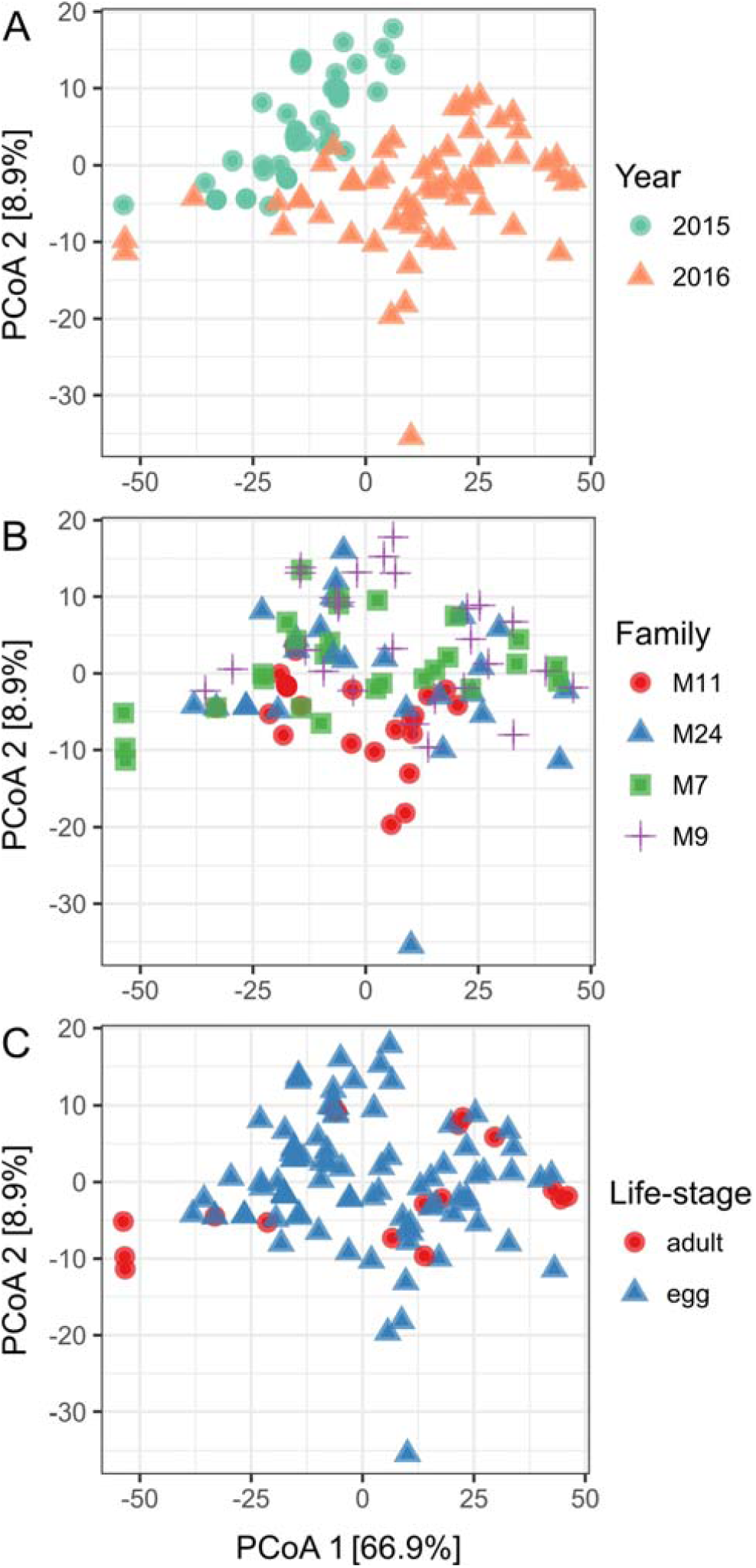
Principal coordinates analysis (PCoA) of log-transformed Manhattan distances among normalized variant counts colored by (A) Sampling year, (B) Family and (C) Life-stage.

A series of linear mixed model analyses further supported the dominant effect of year, with twelve of these variants found to be significantly differentially abundant by year, life-stage or the interaction of the two (figure 1B, 4). Only a single ITS2 sequence variant (sq2) was identified as being significantly differentially abundant between life-stages alone. Sq2, which comprised 17% of the read data across samples, was 1.25-fold more abundant in eggs than adults (P_adj_ < 0.05, figure 4C). Sq4 (5% of reads) was also more abundant in eggs than adults, by 1.27-fold (P_adj_ < 0.05), but showed an additional effect of sampling year, being 1.35-fold more abundant in 2016 than 2015 (P_adj_ < 0.05, figure 4D). Both sq2 and sq4 exhibited a best match to the GeoSymbio Clade C15 reference (figure 4A), and their abundances were marginally positively correlated with one another across samples (R = 0.29). However, they both exhibited strong negative or neutral relationships with the other C15-type variants, suggesting that in spite of sequence similarity, they may not all be intracellular ITS2-type variants (figure S3).

Abundances of ten sequence variants differed between sampling years, including sq4. Eight sequence variants, all C-15 types, comprising 6.6% of total reads, were more abundant in 2016 (bleaching year) than in 2015 (P_adj_ < 0.05, figure 4B, D, F-M). Although the abundance of sq8 increased across sampling years, increases were more pronounced in adults (P_year × life stage_<0.05; figure 4F). In addition, abundances of sq3, sq8, sq10, sq11 and sq20 were generally strongly positively correlated across samples, suggesting that they may represent a cluster of intracellular variants. The largely independent positive relationship found between sq18 and sq21 (R = 0.72) may indicate a separate intracellular type grouping (figure S3). The tenth, a D1-type comprising only 0.04% of the total read data, was significantly less abundant in 2016, by 5.25-fold on average, although this pattern was primarily driven by decreased abundance in eggs and a concomitant tendency for increased abundance in adults (Padj = 0.014; figure 4N). Only one variant, sq5, showed a pattern whereby abundance increased in adults across sampling years but decreased in eggs, such that abundances in the two life stages converged in 2016 (P_adj:year × life stage_ < 0.05, figure 4E).

**Figure 4.**
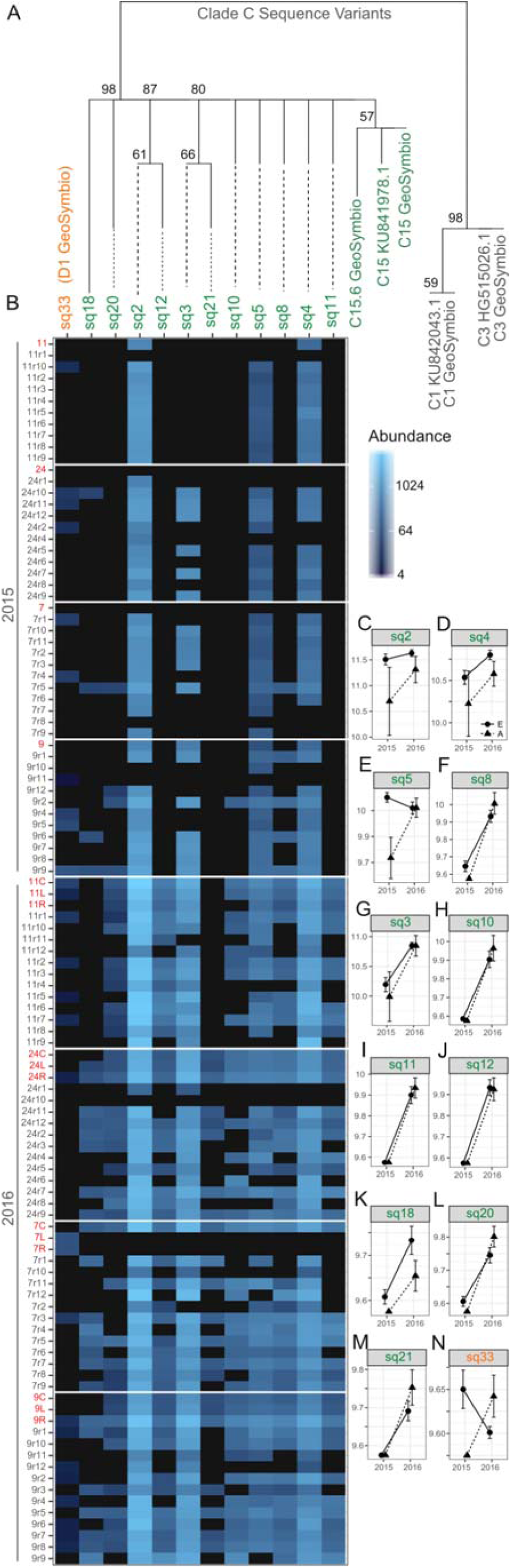
(A) Phylogenetic relationships among significantly differentially abundant (P_adj_ < 0.05) clade C-type OTUs. Reference types are either derived from the GeoSymbio database or from NCBI’s nr database. (B) Heatmap of showing read abundance by sequence variants and samples, grouped by family and ordered by year and life-stage (adult samples are colored red). (C-N) Mean variant abundance ± SEM for statistically significant factors: (C, D) P_life-stage_<0.05, E=eggs, A=adults; (E,F) P_interaction_<0.05; (D,F-N) P_year_<0.05.

### Physiological changes associated with shuffling

Colonies that experienced no bleaching (11, 24) exhibited the greatest relative change in their *Symbiodinium* communities (shuffling score), whereas *Symbiodinium* communities in colonies that bleached (7, 9, bleaching scores 4.8-5.8) remained comparatively constant (figure 5A). Although the sample size was small (n = 4 colonies), a moderate correlation between community change and bleaching status was detected (R^2^ = 0.51, figure 5B).

**Figure 5.**
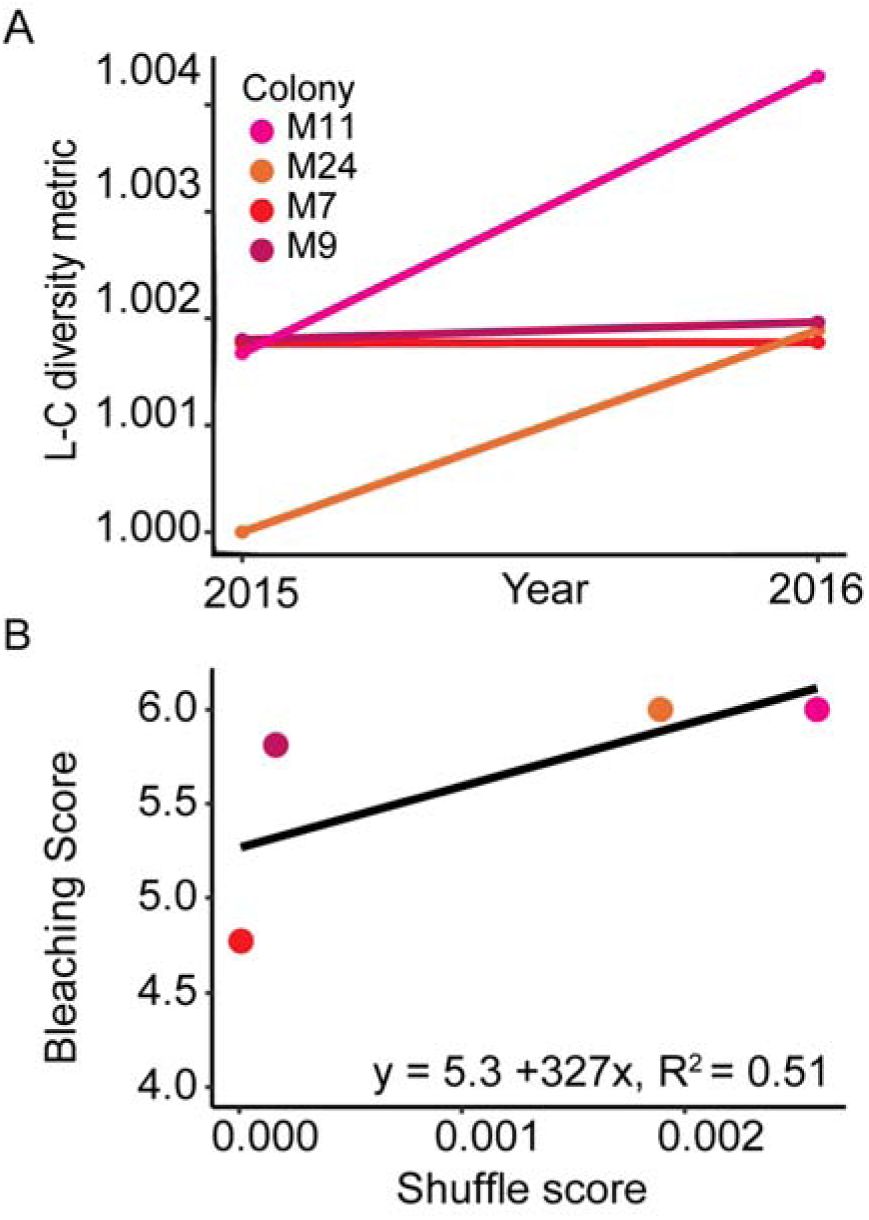
(A) *Symbiodinium* community diversity metric in *M. digitata* during non-bleaching (2015) and bleaching (2016) years for each of the four spawning colonies. (B) Relationship between colony bleaching scores and the magnitude of shuffling in each spawning colony. Low shuffling scores are indicative of communities that did not change between 2015 and 2016, whereas larger shuffling scores are indicative of *Symbiodinium* communities that did change.

## Discussion

This study investigated whether changes in the abundance of *Symbiodinium* types within the endosymbiotic communities of corals in response to a bleaching event are heritable in a coral species capable of vertically-transmitting symbionts to its offspring. Although *Symbiodinium* communities in both adults and eggs of the coral *Montipora digitata* were dominated by one, temporally-stable C15 type (66% of read data), variation in the background or low-abundance *Symbiodinium* community occurred across years. Of the 34 sequence variants identified with high confidence, ten showed significant differences in abundance between a bleaching and non-bleaching year, differences that were consistent in adults and their corresponding eggs, supporting transgenerational inheritance of *Symbiodinium* communities that had been shuffled in parental colonies in response to thermal stress (figure 1, H_2_). In contrast, the abundances of four sequence variants differed among life stages, but patterns were stable across years (H_1_). In two cases, the abundances of sequence variants in eggs changed across years, but abundances differed independently from communities in adults (H3). Overall, the most common pattern in background types was broad mirroring of shuffled adult communities in eggs of *M. digitata.* Our results suggest that not only is the symbiont community composition plastic in adults of *M. digitata,* but also that this plasticity is heritable.

The importance of shuffling background symbiont types to the health of corals is only beginning to be understood. Although some adult corals can shuffle the abundances of *Symbiodinium* types within their endosymbiotic communities in response to temperature changes, thereby increasing the likelihood of survival [48–50], the implications of transgenerational inheritance of these shuffled communities is completely unexplored. In other maternally-inherited symbioses (predominantly bacterial), transmission efficiency of symbionts is influenced by changes in temperature [51,52]. Shuffling and switching of symbionts (secondary acquisition and replacement, respectively) are both prevalent in insects [53], and the acquisition of particular symbionts during temperature stress have been shown to provide protective benefits to the holobiont [54]. Furthermore, symbiont-mediated transgenerational effects have been reported to play a role in immune priming in insects [55] and may be common in plants [56]. Finally, variation in *Symbiodinium* community composition in coral early life-history stages may serve no functional role in vertical transmitters under non-bleaching conditions (e.g. growth, [21], but remains untested under thermal stress. To our knowledge, we present the first evidence that shuffled proportions of symbiont communities in adult corals are largely preserved in a species which transmits *Symbiodinium* to its eggs.

Building on previous evidence of the heritability of symbiont transmission in *M. digitata* [17], our results expand current understanding of the flexibility of the *Symbiodinium* mutualism in a vertical transmitter by showing that these species may have greater flexibility to vary their symbiont communities during times of stress than was previously thought. Early longitudinal studies examining *Symbiodinium* community composition in vertically-transmitting corals in response to naturally occurring thermal stress events led to the general conclusion that communities were highly stable [57,58] although variation in communities across temperature gradients has been observed [59]. Even when some shuffling was observed among adult colonies, communities eventually reverted to their pre-bleaching abundances following a recovery period [13,60,61], suggesting that plasticity was temporally limited, although the length of this recovery period might span years (see [13,24]). Compared to *M. digitata,* congeneric adult colonies of *M. capitata* exhibited no interannual changes of symbiont community composition within individual adult corals or their gametes over time [20]. Although the prevalence of shuffling as an environmental response mechanism is still debated, inter and intraspecific variation has been reported in multiple studies [26,62,63]. Hence, the stability in *M. capitata* can potentially be attributed to the lack of genetic or environmental variation (i.e. non-bleaching conditions insufficient to elicit shuffling) or shallow sequencing depth. Thresholds for shuffling responses may therefore only be brought on by extremely high or fast changes in environmental gradients (e.g. temperature, wave exposure, turbidity; [64]. Regardless of the ultimate temporal persistence of such changes in adult corals, we show here that this altered symbiont community can be transgenerationally inherited, although further work is needed to determine if shuffling thresholds for heritable parental effects are similar to those proposed for adults [26].

Patterns in the changing abundances of background *Symbiodinium* types in response to thermal stress found here for *M. digitata* highlight the complexity in the physiological properties conferred by endosymbiotic *Symbiodinium* on their coral hosts. In horizontal transmitters, the primary pattern described in response to thermal stress so far has been shuffling to increase the abundance of *Symbiodinium* types belonging to clade D (D1 or D1a) during bleaching, followed by a decrease in these types once bleaching conditions subside [11,14,26]. Indeed, shifts to a D-dominated community in adult colonies of *A. millepora* were found to increase thermal tolerance by ~1-1.5°C [12]. While we observed a trend for D1 to increase in adult corals, consistent with the increase in thermal stress across years, this sequence variant comprised less than 0.05% of the overall community. Similarly, nine sequence variants that were significantly more abundant during the 2016 bleaching year in both adults and their eggs, exhibited a best match to a C15 reference type and comprised 12% of the overall community. It has been shown that populations of *Symbiodinium* with the same ITS2-type can exhibit different thermal tolerances, and in symbiosis can also differentially impact physiological limits of the holobiont [8,65]. Correspondingly, the response of *M. digitata* colonies to thermal stress by shuffling to increase the abundance of these different strains of C15 may provide different physiological costs and benefits to their *M. digitata* hosts which may underpin the differences in abundance. Furthermore, differences in the abundances of two sequence variants, a C15 and a D1-type, which increased in adults but did not show similar patterns in eggs, may reflect differences in costs and benefits conferred to hosts between life-stages. Prior work in horizontally-transmitting species has suggested that D1 may not always have a thermal protective function in the early-life stages of corals or across species [66]. Finally, although read depth was significantly lower for 2015 samples due to the two years being processed on independent sequencing runs, directional changes in abundance and random subsampling to mimic reduced sampling effort both suggest that read depth did not alter the main results. Additional information is therefore needed to elucidate the roles that C15 and D1 might have in the *M. digitata-Symbiodinium* partnership.

The potential importance of numerically rare or background symbionts for the coral-algal symbiosis has long been appreciated [67], but appropriate methods for detecting the rare biosphere have only recently become available [24,68,69]. Consequently, the physiological implications of such shifts are only beginning to be explored. Whilst some background symbionts appear to have no functional role [70], recent network analyses of 46 coral genera provide evidence of the importance of cryptic *Symbiodinium* communities, in which rare symbionts appear to significantly contribute to symbiosis stability [71]. Similarly, the rare bacterial biosphere is important to holobiont physiology for a range of both plant and animal symbioses [29] and may act to acclimatize the holobiont to new conditions, as postulated for corals [11,72]. Our finding that adult colonies that exhibited the largest shifts in *Symbiodinium* communities also experienced the lowest levels of bleaching suggests that shuffling symbionts contributes to buffering host physiology under environmental stress. We also found small but significant differences in photochemical efficiency between spawning and non-spawning colonies, although it is unclear whether the magnitude of these differences are biologically significant. Previous research has presented a link between coral ability to shuffle, buffering of photophysiological performance and spawning success when exposed to cold stress [8]. Lower YII values potentially represent smaller deviations away from photophysiological norms, linked to spawning success. Further work on the long-term physiological and fitness consequences of this heritable plasticity would help to evaluate the potential for this mechanism to facilitate rapid coral acclimation to changing environmental conditions.

## Data Availability

All raw sequencing data has been deposited in the NCBI Sequence Read Archive under Accession number SRPXXX.

## Conflict of Interest Statement

The authors declare no conflict of interest.

## Funding

Funding was provided by the US National Science Foundation to CDK (DBI-1401165) and Australian Research Council to BLW (CE1401000020).

## Acknowledgements

We thank Orpheus Island Research Station for spawning assistance. All samples of *M. digitata* from Orpheus Island were collected under Great Barrier Reef Marine Park Authority permit G10/33312.1 to BLW.

## Authors and Contributors

KQ and CK conceived and designed the experiments, performed the laboratory work, data analysis and figure preparation and wrote the manuscript. BW provided technical expertise, provided reagents and critically revised the draft and approved the final version and are in agreement to be accountable for all aspects of the work.

